# Changes in the urinary proteome of rats after short-term intake of magnesium threonate

**DOI:** 10.1101/2023.09.29.559957

**Authors:** Ziyun Shen, Minhui Yang, Haitong Wang, Yuqing Liu, Youhe Gao

## Abstract

Magnesium is an important mineral in living organisms and has multiple functions in the human body, wherein it plays an important therapeutic and preventive role in a variety of diseases. In the present study, urine samples of rats before and after gavage of magnesium threonate were collected, and the urinary proteome was identified using the LC-MS/MS technique and analyzed using various databases. The results illustrated that the urinary proteome of rats was significantly altered after short-term intake of magnesium supplements and that the differential proteins and the biological functions were related to magnesium. This study innovatively establishes a method to study nutrients from the perspective of urine proteomics. This work demonstrates that the urinary proteome is capable of reflecting the effects of nutrient intake on the organism in a more systematic and comprehensive manner and has the potential to provide clues for clinical nutrition research and practice.

## 1 Introduction

Magnesium (Mg) is an essential electrolyte for living organisms. In 1980, Theodor Günther proposed that magnesium acts as a cofactor in more than 300 enzymatic reactions in the body^[1]^. Currently, the Enzyme Database lists over 600 enzymes in which magnesium ions act as cofactors and an additional 200 enzymes in which magnesium ions act as activators^[2–4]^. It is essential for adenosine triphosphate (ATP) metabolism, DNA and RNA synthesis, and protein synthesis^[5]^. In addition, it plays a key role in the regulation of many physiological functions, including muscle contraction, blood pressure regulation, insulin metabolism, cardiac excitability, vasodilatory tone, neurotransmission and neuromuscular transmission^[6]^.

Based on its many functions in the body, magnesium plays an important role in the prevention and treatment of many diseases. Low magnesium levels have been linked to many chronic and inflammatory diseases, such as Alzheimer’s disease, asthma, attention deficit hyperactivity disorder (ADHD), insulin resistance, type 2 diabetes, hypertension, cardiovascular disease, migraines, and osteoporosis[7].

Magnesium is the fourth most abundant mineral in the human body, and humans need a regular intake of magnesium to prevent magnesium deficiency. Many studies have demonstrated the beneficial effects of magnesium supplementation. However, since the recommended daily intake (RDI) of magnesium varies, it is difficult to define the exact optimal intake. According to the Dietary Guidelines for Chinese Residents, the recommended nutrient intake (RNI) for magnesium is 330 mg per day, and the tolerable upper intake level (UL) is 700 mg/d^[8]^. According to the U.S. Food and Nutrition Board, the recommended daily intake of magnesium for adult males and females is 420 mg and 320 mg, respectively^[4]^. The richest source of bioavailable magnesium is the hydrosphere. In seawater, the concentration of magnesium is approximately 55 mmol/liter. Approximately 10% of daily magnesium intake comes from drinking water^[9]^; green vegetables, nuts, seeds, and unprocessed grains are rich sources of magnesium; in addition, some magnesium is found in fruits, fish, meat, and dairy products^[5]^.

Assessing magnesium status in the body is difficult because most magnesium is located intracellularly or in bone. The most common test used in clinical medicine to rapidly assess changes in magnesium status is the serum magnesium concentration, but there is little correlation between serum levels and systemic magnesium levels or concentrations in specific tissues^[10,11]^.

Since urine is not part of the internal environment, in contrast to plasma, there is no homeostasis mechanism in urine, which is thus able to accumulate early changes in the physiological state of the organism and to reflect more sensitively the changes in the organism, making it a good source of biomarkers^[12]^.

In this study, we established a rat model of short-term magnesium threonate intake to investigate the changes in the urinary proteome after short-term supplementation with magnesium supplements to investigate the overall effects of the intake of appropriate amounts of magnesium ions on the organism. This study innovatively establishes a method to study the effect of nutrients on the organism from the perspective of the urine proteome, which can reflect the role of the organism in a more comprehensive and systematic way and can hopefully provide information and clues about nutrient assessment, monitoring and prescription for clinical nutrition research and practice.

## 2 Materials and Methods

### 2.1 Experimental materials

#### 2.1.1 Experimental consumables

Five-milliliter sterile syringes (BD), gavage needles (16-gauge, 80 mm, curved needle), 1.5-ml/2-ml centrifuge tubes (Axygen, USA), 50-ml/15-ml centrifuge tubes (Corning, USA), 96-well cell culture plates (Corning, USA), 10 kD filters (Pall, USA), Oasis HLB solid phase extraction column (Waters, USA), 1-ml/200-μl/20-μl pipette tips (Axygen, USA), a BCA kit (Thermo Fisher Scientific, USA), high pH reverse peptide separation kit (Thermo Fisher Scientific, USA), and iRT (indexed retention time, BioGnosis, UK) were used.

#### 2.1.2 Experimental apparatus

A rat metabolic cage (Beijing Jiayuan Xingye Science and Technology Co., Ltd.), frozen high-speed centrifuge (Thermo Fisher Scientific, USA), vacuum concentrator (Thermo Fisher Scientific, USA), DK-S22 electric thermostatic water bath (Shanghai Jinghong Experimental Equipment Co., Ltd.), full-wavelength multifunctional enzyme labeling instrument (BMG Labtech, Germany), oscillator (Thermo Fisher Scientific, USA), TS100 constant temperature mixer (Hangzhou Ruicheng Instruments Co. BMG Labtech), electronic balance (METTLER TOLEDO, Switzerland), -80 [ultralow-temperature freezing refrigerator (Thermo Fisher Scientific, USA), EASY-nLC1200 Ultra High Performance Liquid Chromatography system (Thermo Fisher Scientific, USA), and Orbitrap Fusion Lumos Tribird Mass Spectrometer (Thermo Fisher Scientific, USA) were used.

#### 2.1.3 Experimental reagents

Magnesium threonate (Shanghai Yuanye Biotechnology Co., Ltd.), Trypsin Golden (Promega, USA), dithiothreitol (DTT, Sigma, Germany), iodoacetamide (IAA, Sigma, Germany), ammonium bicarbonate NH4HCO3 (Sigma, Germany), urea (Sigma, Germany), pure water (China Wahaha Company), mass spectrometry grade methanol (Thermo Fisher Scientific, USA), mass spectrometry grade acetonitrile (Thermo Fisher Scientific, USA), mass spectrometry grade purified water (Thermo Fisher Scientific, USA), Tris-base (Promega, USA), thiosulfate (Sigma, Germany), thiamine (Sigma, Germany), thiophene (Promega, USA), and thiourea (Sigma, Germany) were obtained.

#### 2.1.4 Analysis software

Proteome Discoverer (Version 2.1, Thermo Fisher Scientific, USA), Spectronaut Pulsar (Biognosys, UK), Ingenuity Pathway Analysis (Qiagen, Germany), R studio (Version 1.2.5001), Xftp 7, and Xshell 7 were utilized.

### 2.2 Experimental Methods

#### 2.2.1 Animal modeling

In this study, 14-week-old rats were used to minimize the effects of growth and development during gavage. Five healthy Sprague–Dawley (SD) 9-week-old male rats (250±20 g) were purchased from Beijing Viton Lihua Laboratory Animal Technology Co. The rats were kept in a standard environment (room temperature (22±2)°C, humidity 65%-70%) for five weeks before starting the experiments, and all experimental operations followed the review and approval of the Ethics Committee of the School of Life Sciences, Beijing Normal University.

The dose used in this study was 1208 mg/kg/d magnesium threonate and 100 mg/kg/d elemental magnesium. The tolerable upper intake level (UL) refers to the average maximum daily intake of nutrients at a certain physiological stage and for each sex that are free of any side effects and risks to the health of almost all individuals. According to the Dietary Guidelines for Chinese Residents, the recommended nutrient intake (RNI) for magnesium is 330 mg per day, and the tolerable upper intake level (UL) is 700 mg/d^[8]^. The UL in humans is converted to a rat dose of 65 mg/d according to body surface area and body weight. It has been reported that 604 mg/kg/d magnesium threonate, i.e., 50 mg/kg/d elemental magnesium, is the minimum effective dose for memory enhancement in rat experiments. The dose in the present study was twice this minimum effective dose, three times the recommended intake for humans, and 1.5 times the tolerable upper intake level for humans. First, 24.16 g of magnesium threonate was dissolved in 200 ml of sterile water to configure a magnesium threonate solution. Each rat was gavaged with 5 ml of magnesium threonate solution once a day for 6 days. The first day of gavage was recorded as D1, and so on. Sampling time points were set before and after gavage so that each animal served as its own control. The sample collected on the day before gavage was the control group, recorded as D0, and the sample collected on the 6th day of gavage was the experimental group, recorded as D6.

#### 2.2.2 Urine sample collection

One day before the start of magnesium threonate gavage (D0) and 6 days after magnesium threonate gavage (D6), each rat was individually placed in a metabolic cage at the same time of day, fasted and dehydrated for 12 h. Urine was collected overnight, and the urine samples were collected and placed in the freezer at -80 °C for temporary storage.

#### 2.2.3 Urine sample processing

Two milliliters of urine sample was removed, thawed, and centrifuged at 4 °C and 12,000×g for 30 minutes. The supernatant was removed, and 1 M dithiothreitol (DTT, Sigma) storage solution (40 μl) was added to reach the working concentration of DTT (20 mM). The solution was mixed well and then heated in a metal bath at 37 °C for 60 minutes and allowed to cool to room temperature. Then, iodoacetamide (IAA, Sigma) storage solution (100 μl) was added to reach the working concentration of IAM, mixed well and then reacted for 45 minutes at room temperature protected from light. At the end of the reaction, the samples were transferred to new centrifuge tubes, mixed thoroughly with three times the volume of precooled anhydrous ethanol, and placed in a freezer at -20 °C for 24 h to precipitate the proteins. At the end of precipitation, the sample was centrifuged at 4 °C for 30 minutes at 10,000×g, the supernatant was discarded, the protein precipitate was dried, and 200 μl of 20 mM Tris solution was added to the protein precipitate to reconstitute it. After centrifugation, the supernatant was retained, and the protein concentration was determined by the Bradford method. Using the filter-assisted sample preparation (FASP) method, urinary protein extracts were added to the filter membrane of a 10-kD ultrafiltration tube (Pall, Port Washington, NY, USA) and washed three times with 20 mM Tris solution. The protein was resolubilized by the addition of 30 mM Tris solution, and the protein was added in a proportional manner (urinary protein:trypsin = 50:1) to each sample. Trypsin (Trypsin Gold, Mass Spec Grade, Promega, Fitchburg, WI, USA) was used to digest proteins at 37 °C for 16 h. The digested filtrate was the peptide mixture. The collected peptide mixture was desalted by an Oasis HLB solid phase extraction column, dried under vacuum, and stored at -80 °C. The peptide mixture was then extracted with a 0.1% peptide mixture. The lyophilized peptide powder was redissolved by adding 30 μL of 0.1% formic acid water, and then the peptide concentration was determined by using the BCA kit. The peptide concentration was diluted to 0.5 μg/μL, and 4 μL of each sample was removed as the mixed sample.

#### 2.2.4 LC-MS/MS tandem mass spectrometry analysis

All identification samples were added to a 100-fold dilution of iRT standard solution at a ratio of 20:1 sample:iRT, and the retention times were standardized. Data-independent acquisition (DIA) was performed on all samples, and each sample measurement was repeated 3 times, with 1-mix samples inserted after every 10 runs as a quality control. The 1-μg samples were separated using EASY-nLC1200 liquid chromatography (elution time: 90 min, gradient: mobile phase A: 0.1% formic acid, mobile phase B: 80% acetonitrile), the eluted peptides were entered into the Orbitrap Fusion Lumos Tribird mass spectrometer for analysis, and the corresponding raw files of the samples were generated.

#### 2.2.5 Data processing and analysis

The raw files collected in DIA mode were imported into Spectronaut software for analysis, and the highly reliable protein criterion was a peptide q value<0.01. The peak area quantification method was applied to quantify the proteins by applying the peak area of all fragment ion peaks of the secondary peptides.

Proteins containing two or more specific peptides were retained, and the contents of individual proteins identified in different groups of samples were compared to screen for differential proteins. The conditions for screening differential proteins were a p value <0.05 for paired t test analysis and fold change (FC) >5 or <0.2.

Unsupervised cluster analysis, such as hierarchical cluster analysis (HCA) and principal component analysis (PCA), was performed using the wkomics platform (https://omicsolution.org/wkomics/main/), and functional enrichment of differential proteins was performed using the Database for Annotation, Visualization and Integrated Discovery (DAVID) (https://david.ncifcrf.gov/). Gene Ontology (GO) analysis was conducted to obtain the results from the 3 aspects of biological process, cellular localization and molecular function. Ingenuity Pathway Analysis (IPA) was performed for the identified differential proteins, and a search was conducted for the differential proteins and related pathways based on the PubMed database (https://pubmed.ncbi.nlm.nih.gov/).

#### 2.2.6 Randomized grouping analysis

A total of 10 samples before (n=5) and after (n=5) gavage were randomly divided into two groups, and the mean of the number of differential proteins in all randomized combinations was calculated according to the screening conditions of P value <0.05 and fold change (FC) >5 or <0.2.

## 3 Results and discussion

### 3.1 Analysis of urinary proteome changes before and after ingestion of magnesium threonate

A total of 1766 proteins were identified in all samples (meeting unique peptides>1 and FDR<1%). The results of unsupervised cluster analysis and principal component analysis (PCA) showed that the two groups of samples could be well distinguished, indicating significant changes in the rat urine proteome after continuous gavage of magnesium threonate for 6 days, as shown in Figures 1 and 2.

**Figure 1.**
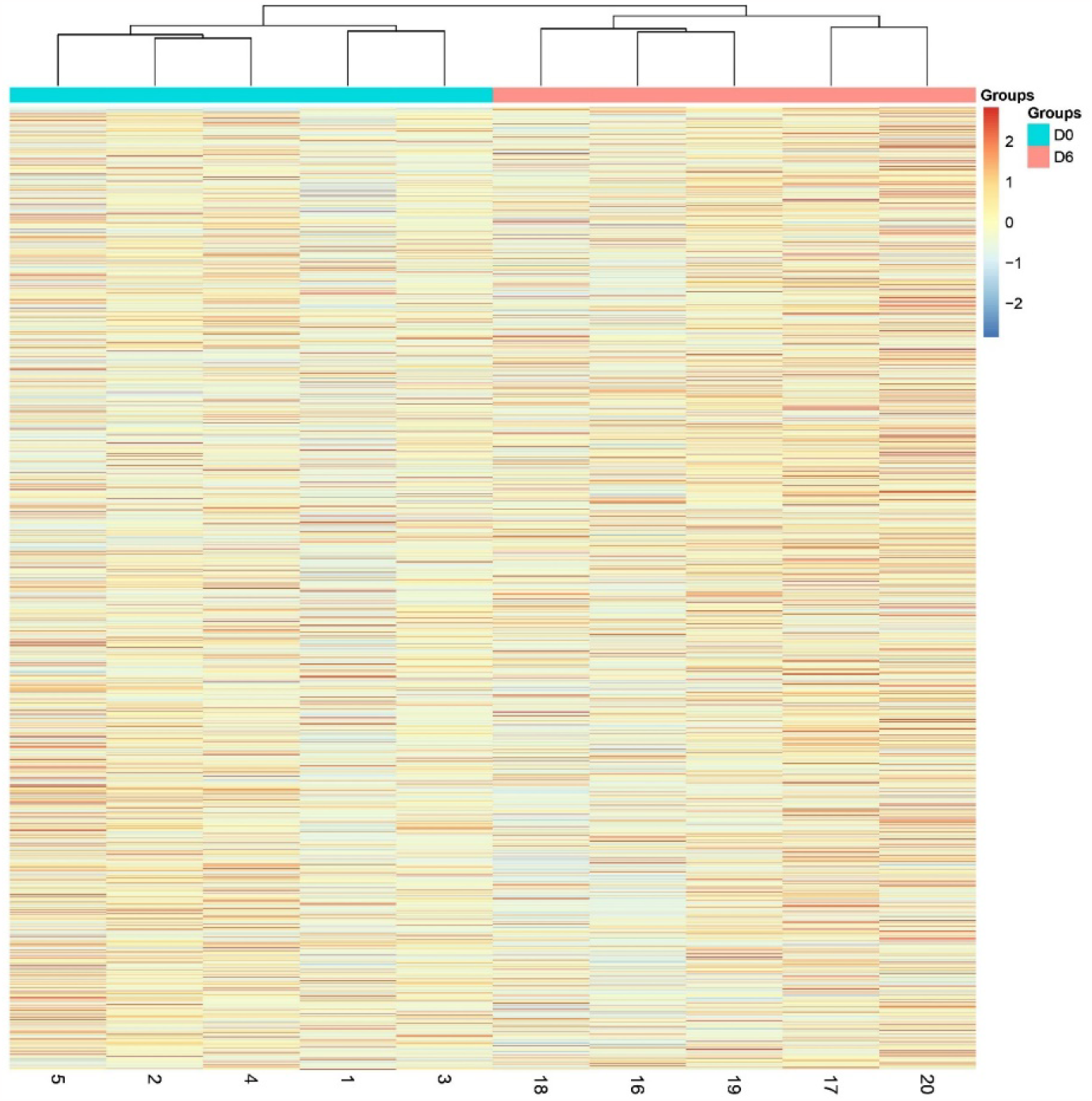
Hierarchical cluster analysis (HCA) of the total proteins in the samples

**Figure 2.**
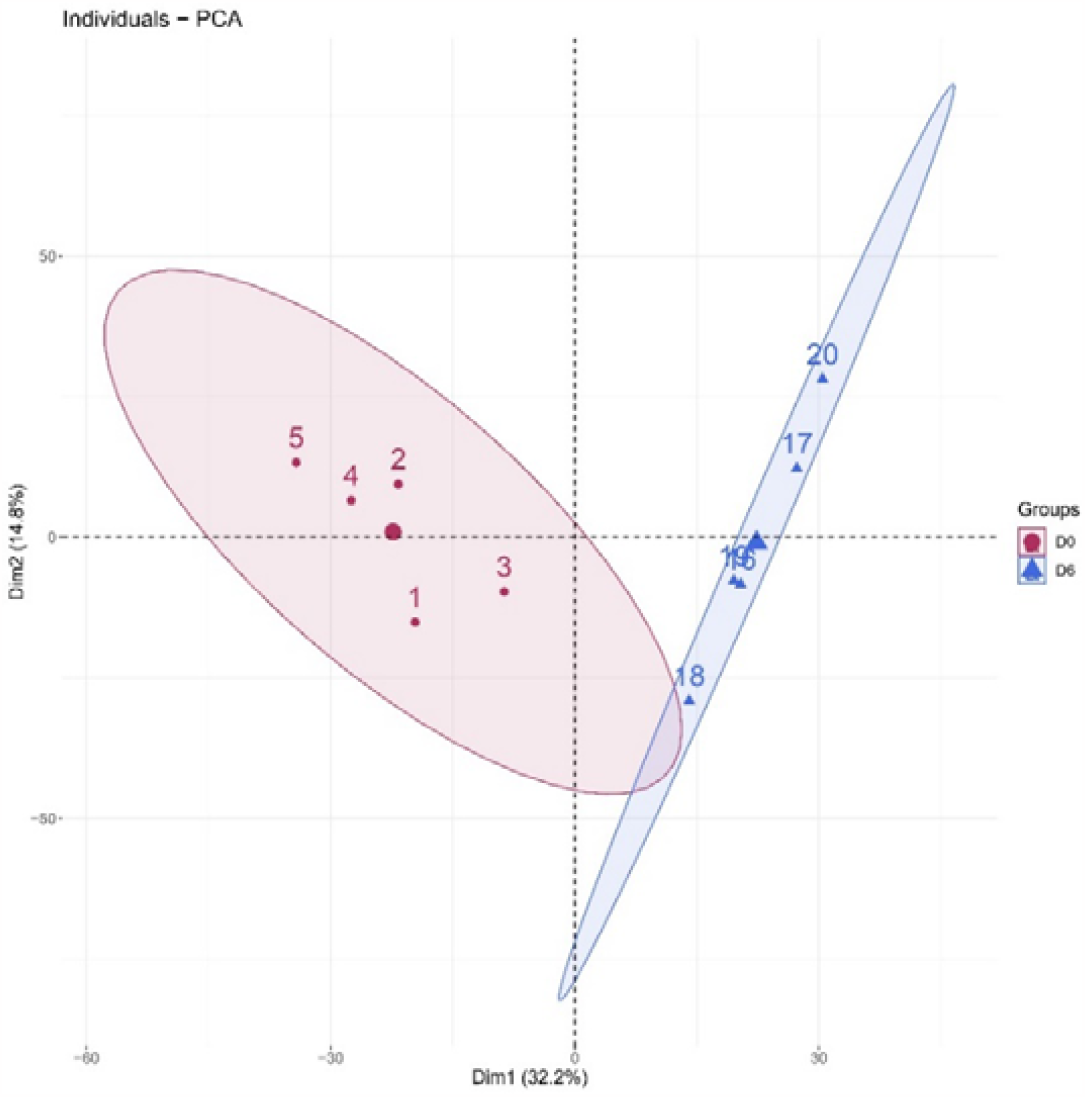
Principal component analysis (PCA) of the total proteins in the samples

### 3.2 Randomized grouping test results

We used a randomized grouping statistical analysis strategy, which is suitable for the study of proteomes with limited sample sizes and for determining whether the differences between two groups are randomly generated^[13]^.

To determine the likelihood of random generation of the identified differential proteins, we validated the random grouping of total proteins identified from two groups of 10 samples, applying the same criteria for the screening of differential proteins, that is, FC ≥ 5 or ≤ 0.2 and P < 0.05, and performing 126 random groupings yielded an average of 20.53 differential proteins, with a proportion of randomly identified proteins of 8.18%, indicating that at least 91.82% of the differential proteins were not due to randomness. The results of the randomized grouping test are shown in Table 1. The probability that the 251 differential proteins we identified by screening were randomly generated is low, and the results show that these differential proteins are indeed associated with short-term intake of magnesium threonate supplements.

**Table 1.**
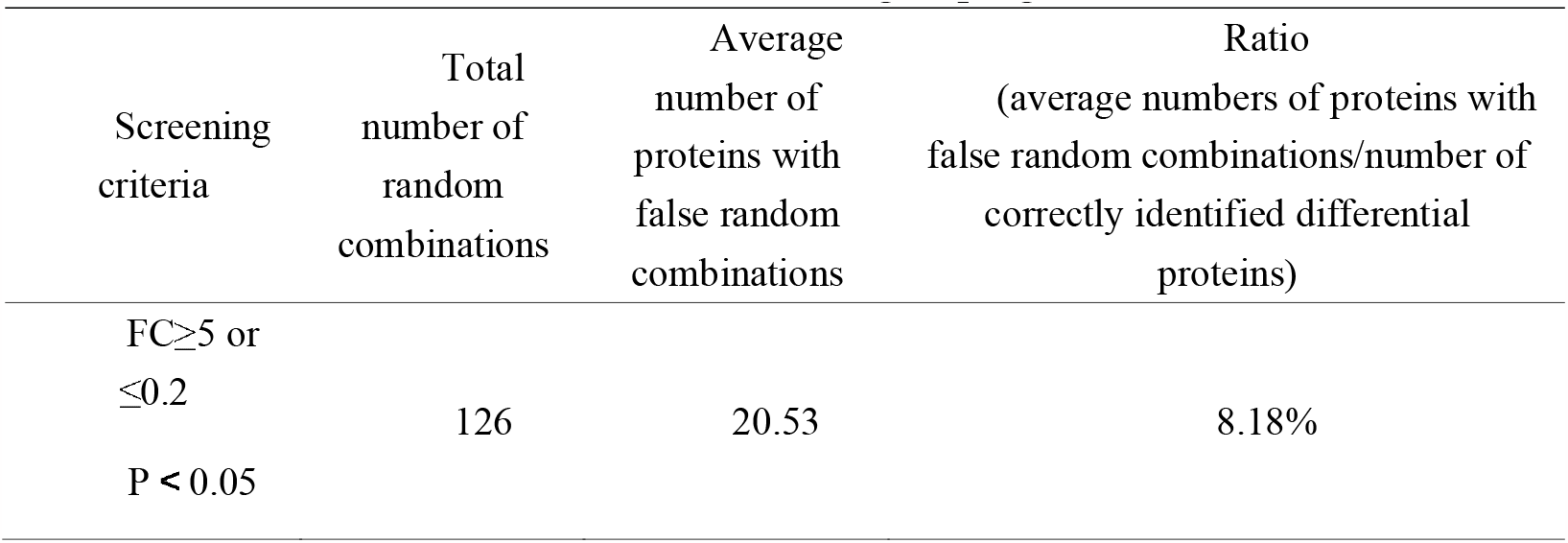
Randomized grouping results.

### 3.3 Differentially expressed protein function annotation

The missing values were replaced with 0. The pregavage samples of rats with magnesium threonate were compared with the samples on Day 6 of gavage, and 251 differential proteins were identified under the screening conditions of FC ≥ 5 or ≤ 0.2, and P < 0.05, as detailed in the supplementary table.

Among them, 63 differential proteins showed significant changes with the largest fold changes (FC=0 or ∞), such as solute carrier family 21 (SLC21), angiotensin-converting enzyme, collagen type X alpha 1 chain, and heat shock 70 kDa. The fold change of 58 differential proteins, including collagen type X alpha 1 chain, heat shock 70 kDa protein, integrin alpha 3, and protein-tyrosine-phosphatase (PTP), was zero, which means that they were detected in the urinary proteome of rats before gavage but were so low as to be nondetectable in the urinary proteome of rats after six days of gavage. Four differential proteins, including PDZ and LIM domain protein 1, had a fold change of ∞, i.e., the protein was undetectable in the rat urine proteome before gavage but was detectable in the rat urine proteome after six days of gavage.

Functional annotation and searching of these differential proteins using the UniProt database and PubMed database revealed that many of the significantly changed differential proteins were directly or indirectly related to magnesium ions, as shown in Table 2.

**Table 2.**
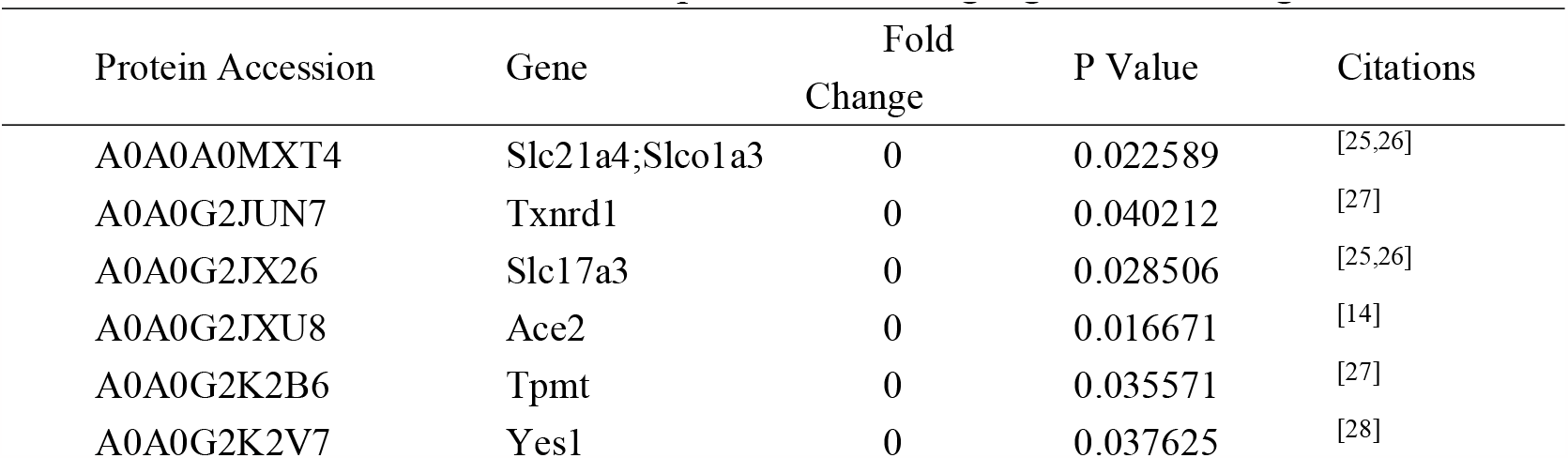
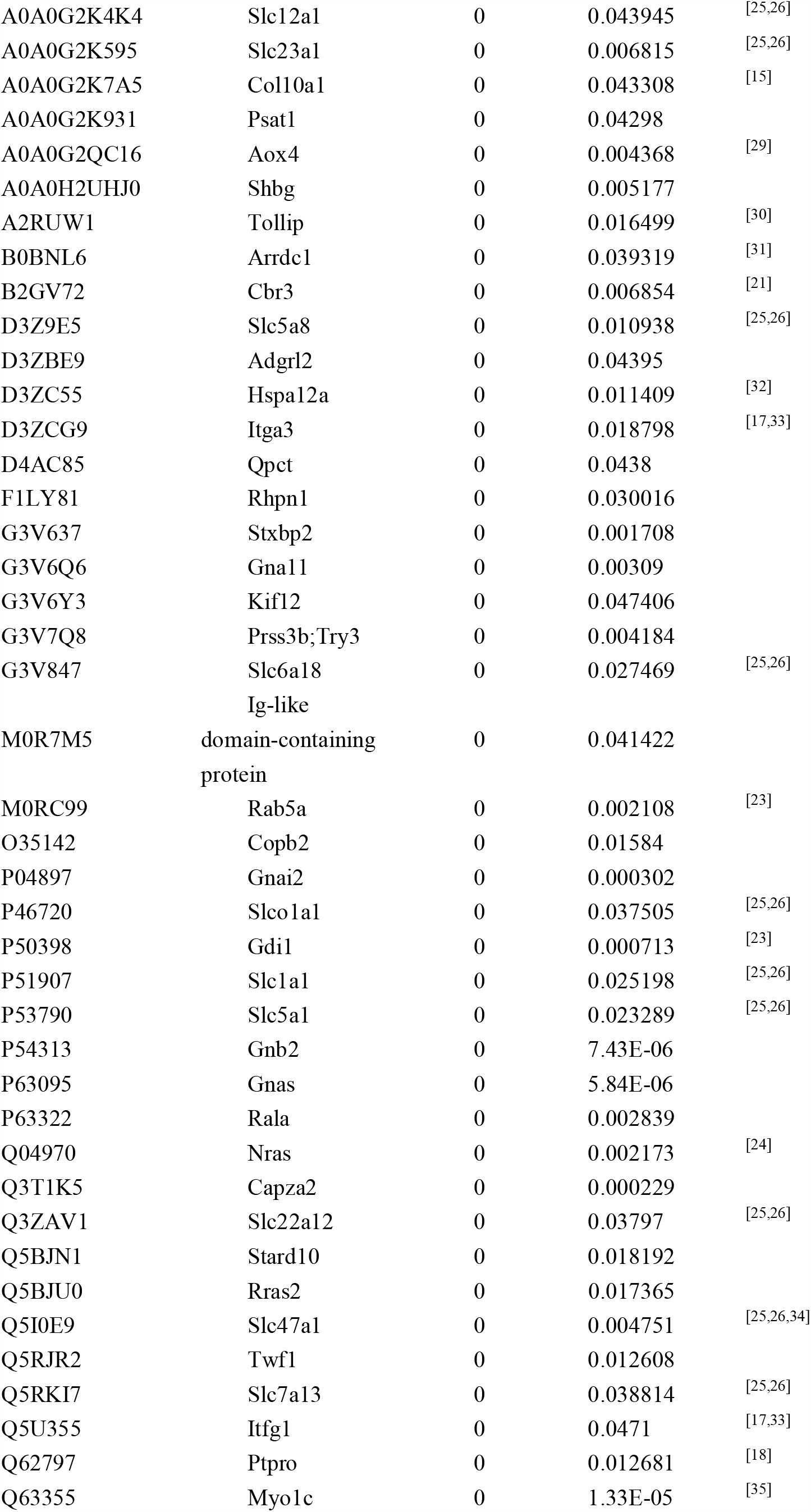
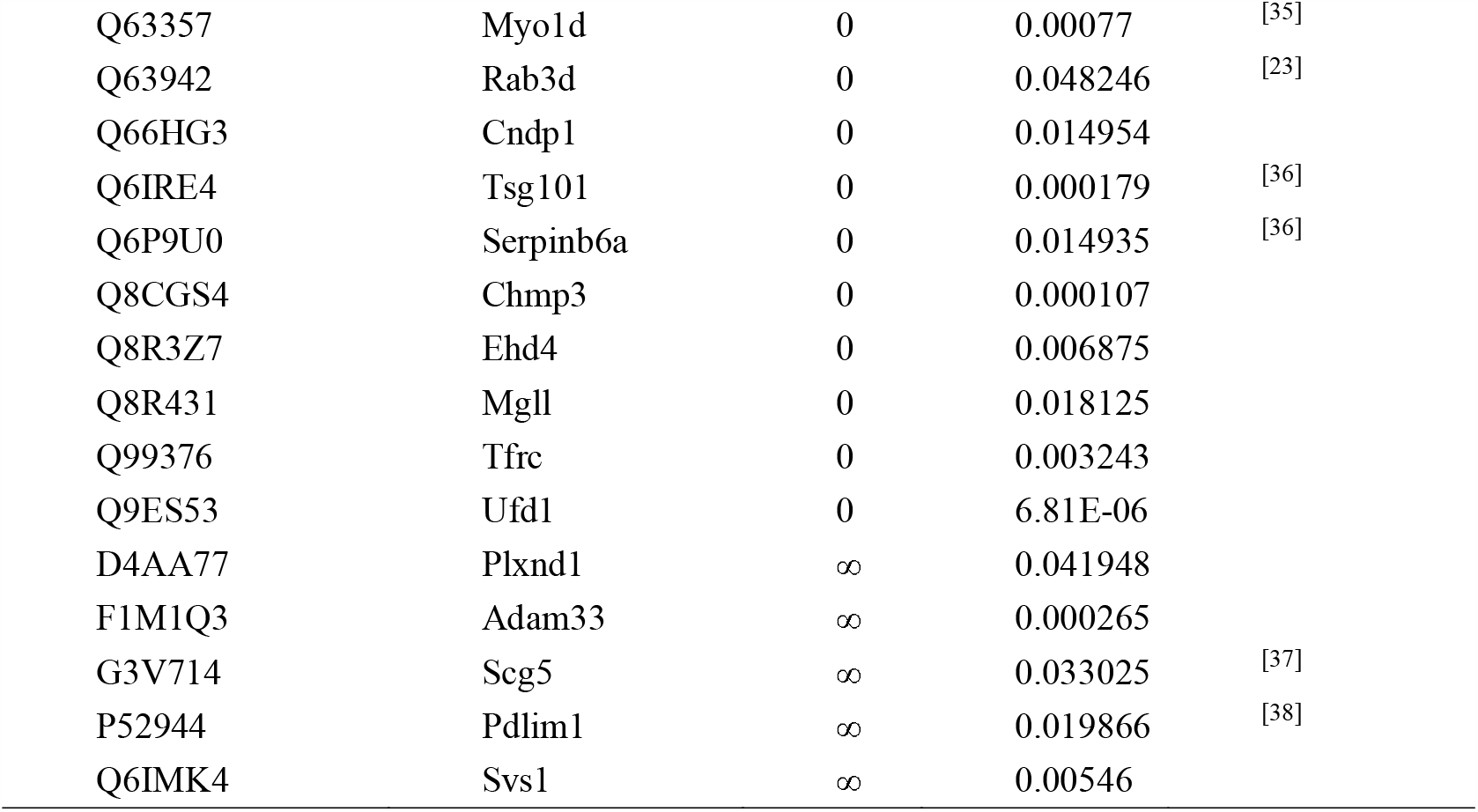
Differential proteins showing significant changes.

According to the literature, the expression levels or activities of many differential proteins are positively or negatively correlated with magnesium ion levels.

Angiotensin-converting enzyme is a peptidase involved in the formation of angiotensin II and the inactivation of bradykinin. In one study, women with gestational hypertension had elevated angiotensin-converting enzyme activity before treatment with magnesium sulfate and lower levels during treatment[14]. Magnesium ion intake may influence blood pressure regulatory mechanisms.

In in vitro culture experiments, Mg^2+^ promoted bone regeneration by promoting osteoblast type X collagen and VEGF production in bone tissue^[15]^.

Magnesium supplementation inhibits the upregulated expression of HSP70 and HSP70 mRNA in vivo^[16]^.

Magnesium ions promote cell proliferation mediated by integrin α2 and integrin α3^[17]^. Magnesium and zinc ions activate and inhibit protein tyrosine phosphatase 1B (PTP1B), respectively^[18]^. Magnesium inhibits the collagen-cleaving activity of the 41-kDa serine-dependent protease, and calcium protects the protease from this inhibition^[19]^. Magnesium sulfate inhibits LPS-macrophage binding by decreasing CD14 expression, and these mechanisms may involve the activation of serine proteases^[20]^.

Many processes in which differential proteins function require the involvement of magnesium ions. For example, magnesium ions play an important role in NADPH oxidase function^[21]^. Of the differential proteins showing significant changes with or without magnesium ions, at least five differential proteins were associated with ATPase and ADP/ATP binding, and at least five were associated with GTPase and GDP/GTP binding. Magnesium ions have the ability to form chelates with ATP and GTP, and many ATPases and GTPases are magnesium-dependent enzymes^[22]^. In addition, Rab5a and NRAS were two differential proteins that exhibited significant changes. Mg^2+^ ions interact with the GTP form and GDP form of Rab5 and Rab7^[23]^. Magnesium ions play a bridging role in the interaction of ligands with NRAS, favoring the association of GDP, GTP and GNP with NRAS^[24]^.

A number of differential proteins have been associated with the transport of magnesium ions. For example, a total of 14 differential proteins belong to the solute carrier family, and the solute carrier family members 1 and 2 (SLC41A1 and SLC41A2) channels of this family are capable of transporting magnesium ions^[25]^.

### 3.4 Molecular functional analysis of differentially expressed protein enrichment

Using the DAVID database to enrich and analyze the molecular functions of 251 differential proteins (screening conditions FC ≥ 5 or ≤ 0.2 and p < 0.05), 45 molecular functions, such as calcium binding, protein binding, insulin-like growth factor binding, integrin binding, GTP binding, PDZ domain binding, transmembrane transport, enzyme regulation, and enzyme activity, were enriched (p < 0.05). The molecular functions, protein numbers and p values are shown in Figure 3.

**Figure 3.**
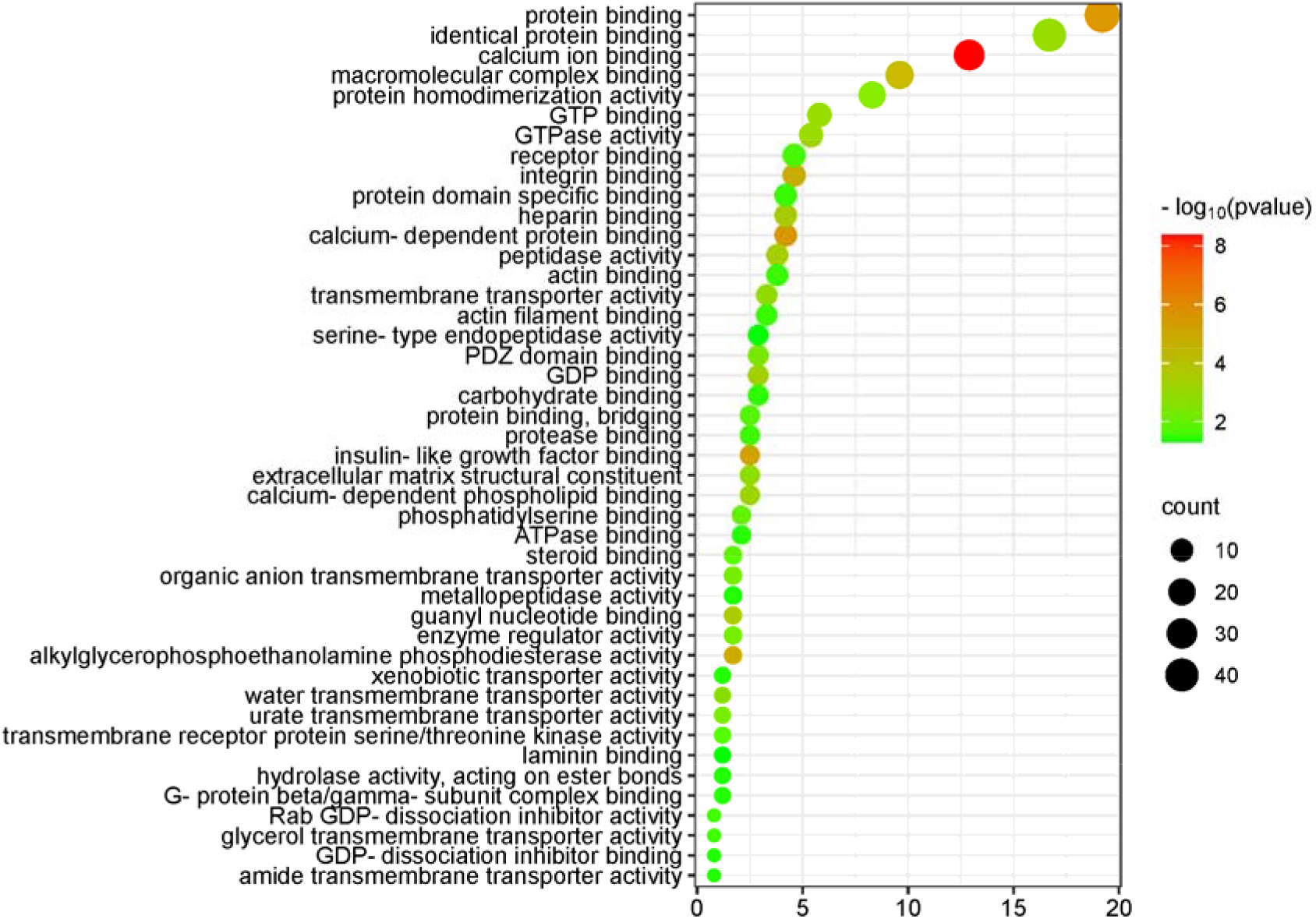
Molecular function (MF) enrichment analysis of the differential proteins (DAVID database, GO analysis)

The five molecular functions with the highest significance were calcium binding, protein binding, calcium-dependent protein binding, insulin-like growth factor binding, and integrin binding.

The molecular functions of calcium ion binding, calcium-dependent protein binding, and calcium-dependent phospholipid binding, which are closely related to calcium ions, were enriched in 35 differential proteins. The molecular function of protein binding was enriched in 46 differential proteins. Magnesium ions have the ability to compete with calcium ions for binding sites on proteins and membranes, and magnesium contributes to the maintenance of a low resting intracellular concentration of free calcium ions, which is important in many cellular functions^[39,40]^.

The molecular function of insulin-like growth factor binding was enriched in six differential proteins. According to the literature, insulin-like growth factor 1 (IGF-1) plays a role in cellular magnesium metabolism and can enhance insulin sensitivity^[41]^. Magnesium exerts insulin sensitization through autophosphorylation of the insulin receptor and regulation of tyrosine kinase activity on the receptor^[42]^. Magnesium is an essential cofactor for several enzymes involved in carbohydrate metabolism^[43]^. Magnesium deficiency is associated with an increased incidence of diabetes mellitus^[44]^.

Integrin binding is a molecular function that was enriched in 11 differential proteins. Magnesium ions promote cell attachment through binding interactions between the integrin family and FAK-related signaling pathways^[33]^.

Magnesium can regulate ion transport via pumps, carriers and channels^[45]^. Many of the molecular functions enriched by differential proteins are related to transmembrane transport, including transmembrane transporter protein activity, water transmembrane transporter protein activity, urate transmembrane transporter protein activity, organic anion transmembrane transporter protein activity, transmembrane receptor protein serine/threonine kinase activity, glycerol transmembrane transporter protein activity, heterotrimeric transporter activity, and amide transmembrane transporter activity, among other molecular functions.

Water transmembrane transporter activity was enriched in three differential proteins, and the fold change of aquaporin-1 (AQP-1), a differentially expressed protein, was 0.005. According to the literature, magnesium ions regulate the expression of the water channel proteins AQP-3 and AQP-4^[46]^.

The molecular function of PDZ domain binding was enriched in eight differential proteins. It has been reported in the literature that PDZ domain containing ring finger 3 (PDZRN3) is able to regulate the magnesium regulator claudin-16 localized in renal tubular epithelial cells^[38]^.

Enzyme-related molecular functions enriched in differential proteins included alkylglycerophosphoethanolamine phosphodiesterase activity, peptidase activity, GTPase activity, enzyme regulatory activity, transmembrane receptor protein serine/threonine kinase activity, protease binding, hydrolase activity, metalloproteinase activity, and serine-type endopeptidase activity.

At least 60 differential proteins belong to the class of enzymes and are closely related to the synthesis of sugars, proteins, nucleic acids and other molecules; signal transduction; and energy metabolism processes.

Mg is a cofactor in hundreds of enzymatic reactions, including those mediated by phosphotransferases and hydrolases such as ATPase^[22]^. Magnesium affects enzyme activity by binding to a ligand, binding to the active site of an enzyme, causing a conformational change in the catalytic process, promoting the aggregation of multienzyme complexes, or other mechanisms^[22]^. Magnesium-dependent enzymes include Na/K-ATPase, hexokinase, creatine kinase, protein kinase, aldehyde dehydrogenase, and cyclases^[40]^. These enzyme systems regulate a variety of biochemical reactions in the body, including protein synthesis, muscle and neurotransmission, neuromuscular transmission, signal transduction, blood glucose control, and blood pressure regulation. Magnesium is required for protein and nucleic acid synthesis, the cell cycle, cytoskeletal and mitochondrial integrity, energy production, and binding of substances to the plasma membrane^[5]^.

Differential proteins enriched molecular functions such as GTP binding, GDP binding, GTPase activity, Rab GDP-dissociation inhibitor activity, G-protein beta/gamma-subunit complex binding, GDP-dissociation inhibitor binding and other molecular functions. Magnesium ions have the ability to form chelates with the important anionic ligand GTP, and GTPases are magnesium-dependent enzymes^[22]^. Magnesium ions play a role in the G protein-coupled receptor signaling pathway^[47]^.

The two molecular functions of actin binding and actin filament binding were enriched in a total of 14 differential proteins. Magnesium regulates actin binding and ADP release in the myosin motor domain^[48]^.

### 3.5 Analysis of the biological processes with differentially expressed protein enrichment

The DAVID database and IPA database were used to enrich and analyze the biological processes and biological pathways of 251 differential proteins (the screening conditions were FC ≥ 5 or ≤ 0.2, and p < 0.05).

The KEGG pathway results for differentially expressed protein enrichment are shown in Figure 4; the biological process (BP) results are shown in Supplementary Figure 1; and the IPA results are shown in Supplementary Figure 2.

**Figure 4.**
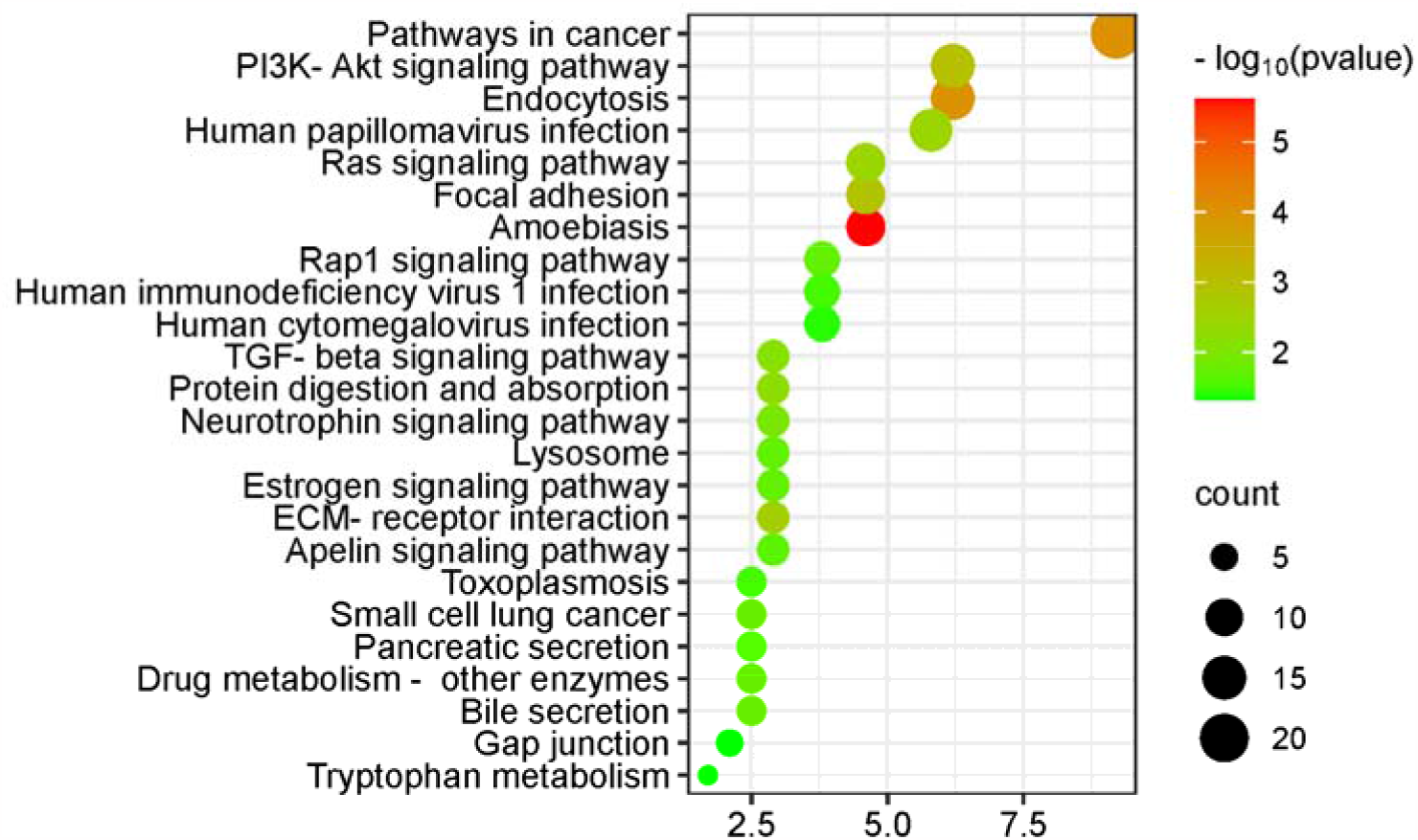
KEGG pathway enrichment analysis of the differential proteins (DAVID database, GO analysis)

The differential proteins enriched many KEGG pathways related to cancer and immunity, including the pathway of cancer, TGF-β signaling pathway, extracellular matrix (ECM) receptor interaction pathway, and small cell lung cancer. Biological processes that were enriched included regulation of cell migration and regulation of cell death. Magnesium ion levels have been reported to be associated with cancer risk and treatment outcome^[49]^.

In addition, magnesium ions have been reported to enhance the transfer of human oncovirus E2 protein from a nonspecific binding site to a specific binding site^[50]^. High levels of magnesium ions can act as a therapeutic agent for chronic amoebiasis^[51]^.

Biological processes such as leukocyte migration and the adaptive immune response are associated with the immune response. Magnesium plays a role in immune function, and magnesium ions are involved in macrophage activation, adherence, and bactericidal activity of the granulocyte oxidative burst; lymphocyte proliferation; and endotoxin binding to monocytes^[6]^.

The PI3K-Akt signaling pathway was enriched in 15 differential proteins. The PI3K-Akt signaling pathway is the classical response to insulin signaling, which is mediated by serine or threonine phosphorylation of a series of downstream substrates. The key genes involved are phosphatidylinositol 3-kinase (PI3K) and AKT/protein kinase b. There is sufficient evidence that magnesium ions activate the PI3K-Akt signaling pathway^[52,53]^.

At the same time, the biological process of regulation of glucose metabolic process was enriched in three differential proteins, and four differential proteins were associated with the positive regulation of insulin secretion, as detailed in the Supplementary Figure 1. According to the latest guidelines from the Association for Magnesium Research, magnesium supplementation can benefit diabetic individuals in different ways: insulin sensitization, calcium antagonism, blood pressure regulation and endothelial stabilization^[54]^.

The neurotrophin signaling pathway, a cell signaling mechanism that plays a key role in the development, plasticity and repair of the nervous system, was enriched in seven differential proteins.

The biological processes which the differential proteins enriched included learning or memory, axon guidance, axon regeneration, positive regulation of neuron projection development, and nervous system development. Magnesium (Mg) is a modulator of glutamatergic N-methyl-D-aspartic acid (NMDA) receptor affinity and can regulate nerve-muscle junction excitability, which may be related to the neurobiological characteristics of mood disorders and psychiatric disorders. Animal experiments have revealed that magnesium can improve hippocampal synaptic plasticity and learning memory in rats^[55]^. Magnesium has the potential to be used as an adjunctive drug in the treatment of depression, anxiety, attention deficit hyperactivity disorder and other psychiatric disorders and Alzheimer’s disease^[37,56]^.

The KEGG pathway was enriched for seven differential proteins in the apelin signaling pathway. The apelin signaling pathway plays an important role in the maintenance of endothelial homeostasis and immunomodulation as well as in the protection of lipids from oxidation, and magnesium sulfate can be used to regulate serum apelin levels^[57]^.

Differential proteins enriched biological processes including angiogenesis, blood coagulation, and heart development. Dietary magnesium deficiency plays an important role in the pathogenesis of ischemic heart disease, congestive heart failure, sudden cardiac death, cardiac arrhythmias, diabetic vascular complications, preeclampsia/eclampsia, and hypertension^[6]^. Several studies have also shown the beneficial effects of magnesium supplementation in the treatment of the above diseases.

The differential proteins enriched many biological processes related to transmembrane transport, including protein transport, organic anion transport, sodium ion transport, extracellular transport, water transport, and responses to various ions, such as mercury, iron, calcium, cadmium, zinc, and copper. Magnesium ions often regulate ion transport through pumps, carriers, and channels^[45]^, which in turn can regulate signaling and the cytoplasmic concentration of ions such as calcium and potassium.

The adenylate cyclase-activating G-protein coupled receptor signaling pathway was enriched in six differential proteins, and the biological process of small GTPase-mediated signaling was enriched in five differential proteins. Magnesium ions have been reported to directly activate adenylate cyclase^[22]^ and to play a role in the G protein-coupled receptor signaling pathway^[47]^.

Differentially expressed protein enrichment in biological processes included bone resorption, osteoblast differentiation, osteoclast proliferation, osteoclast differentiation, ossification, and processes related to bone development and bone metabolism. The literature suggests a causal relationship between dietary intake of magnesium (Mg) and maintenance of normal bone.

The biological process of lung development was enriched for seven differential proteins. The literature suggests that dietary magnesium intake is associated with lung function, airway responsiveness and respiratory symptoms^[58]^.

The estrogen signaling pathway was enriched for seven differential proteins (see Figure 4). Many hormones, including sex steroids, have been reported to affect magnesium homeostasis^[59]^.

## 4 Prospective

Micronutrients (MNs) include minerals and vitamins, and mineral elements can be categorized into macrominerals and microminerals according to human needs. Unlike macronutrients such as carbohydrates, proteins, and fats, micronutrients are required by the human body in very small amounts^[60]^. Although micronutrient requirements are small, micronutrients play an important role in regulating and maintaining various physiological functions in the human body; are essential for cellular function, antioxidant defense and enzymatic reactions; and are involved in many processes, such as immune function, bone health, coagulation, and neurological function^[61,62]^. Several vitamins (including vitamins A, B6, B12, C, D, E, and folic acid) and trace mineral elements (including zinc, iron, selenium, magnesium, and copper) have been reported to play important and complementary roles in supporting the innate and adaptive immune systems. Micronutrient imbalances can negatively impact immune function and can reduce resistance to infection^[63]^. Epidemiologic studies have shown that intake levels of trace minerals such as magnesium, iron, zinc, and copper are associated with the incidence of coronary artery disease^[64]^. Much of the literature suggests that magnesium can play an important therapeutic and preventive role in a wide range of diseases, such as diabetes, osteoporosis, bronchial asthma, preeclampsia, migraines and cardiovascular disease^[6]^, and has an adjunctive effect on the efficacy of certain medications. Deficiencies or excesses of various types of micronutrients are closely associated with various diseases.

It is evident that micronutrients are essential for good health, and deficiencies or excesses of micronutrients may lead to dysfunctions in the body, affecting signaling disorders, cellular processes and organ dysfunctions, which can lead to the development of pathological conditions^[65]^. The impact of nutrient supplementation on the prevention and treatment of diseases should not be ignored, and the study of the role of nutrient intake on the organism is important for disease monitoring, assessment of the effectiveness of treatment and prognosis. Therefore, it is meaningful to explore biomarkers related to nutrient intake.

Previous nutritional studies have usually been limited to focusing on the content or intake of the nutrient itself. The most common method is to measure the concentration of a particular nutrient in a sample of body fluid. However, for some nutrients, extrapolating the level of body stores by directly assessing specific levels of a particular nutrient in body fluids is inaccurate and often invasive. In the case of magnesium ions, for example, assessing magnesium status in the body is difficult because most magnesium is located intracellularly or in bone. The most common method used in clinical medicine to rapidly assess changes in magnesium status is the measurement of serum magnesium concentrations, but serum levels have little correlation with systemic magnesium levels or concentrations in specific tissues^[10]^. Some researchers have also chosen to use metabolites of foods as markers of certain nutrient intakes^[66]^. Some researchers use subjective dietary recall for dietary assessment^[67]^. However, this type of intake assessment is not very accurate and only reflects intake, not the body’s reflection of the nutrient.

To the author’s knowledge, no study has yet reserch the overall effect of nutrients on the organism from a urinary proteomic perspective.

Urinary proteins originate from the body and can reflect small changes in the body and even have the potential to reflect the corresponding biological processes, signaling pathways and molecular mechanisms, which can reflect changes in the physiological state of the organism in a more comprehensive and systematic manner. Our study establishes a novel approach to study nutrients, focusing on the urine proteome, to investigate how short-term supplementation with moderate amounts of nutrients affects the body. It provides a new research perspective in nutrition, which is groundbreaking in the field of nutrition, and has the potential to provide information and clues about nutrient assessment and monitoring for clinical nutrition research and practice.

The results of the study illustrate that the urinary proteome of rats can reveal changes in proteins and biological functions related to magnesium ions after short-term intake of magnesium. In conclusion, short-term supplementation with magnesium can have an effect on the organism, and the urinary proteome reflects this change more comprehensively and systematically.

