## Supplementary figures and images for "Changes in the urinary proteome of rats after short-term intake of magnesium threonate"

### Supplementary Figure 1

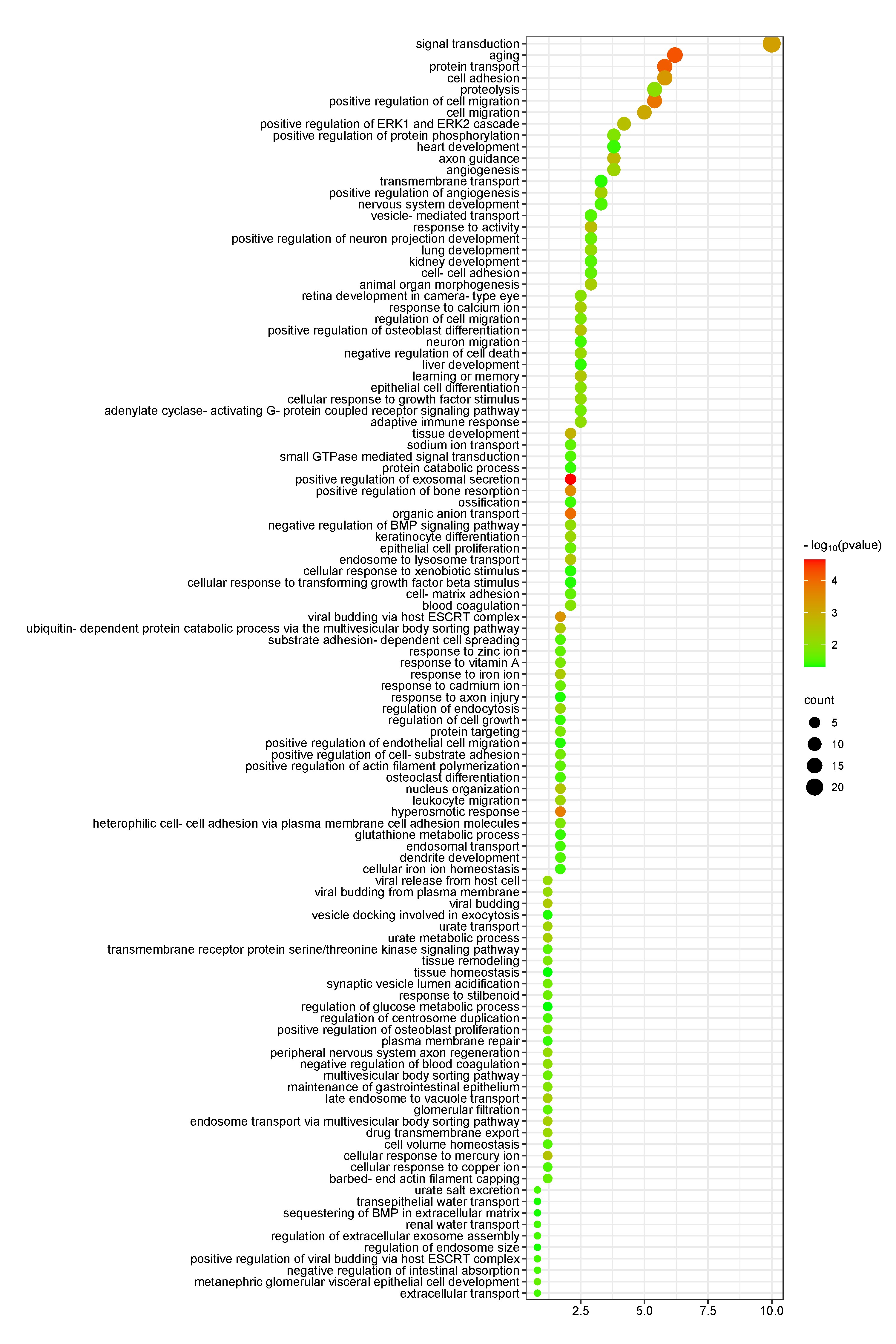

### Supplementary Figure 2

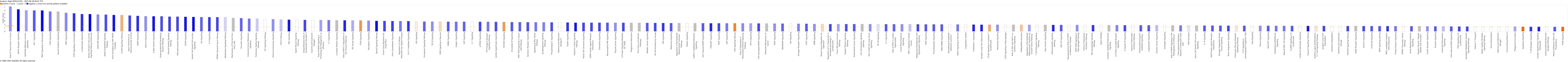
